# Apoptosis-coupled senescence causes cancer cell senotherapy

**DOI:** 10.1101/2023.09.22.558973

**Authors:** Byung-Soo Youn

## Abstract

Although new generations of anti-cancer modalities have been accumulated involving immuno-oncology cancers remain prevailing. This implies the current understanding of cancer cell biology is far from satisfactory. Curation of cancers is extremely rare. We hypothesized what could be the easiest Achilles’ Hill of cancer cells such that simple administration can jab cancer cells to be knocked out. Of conspicuous differences between cancer cells and normal cells, for example, metabolism, hypoxia, anaerobic glycolysis, uncontrolled cell proliferation, etc. exist. What could be the easiest and the most reliable anti-cancer modalities? We discovered one was cancer cell senescence (CCS) because cancer cells are the most presenescent (old) cells. We utilized a synthetic polyphenol designated as ONG41008. ONG41008 was able to induce massive senescence of pathologic myofibroblasts (pMFBs) and a vast majority of representative human cancer cells as well as a line of primary NSCLCs. All these cells turned out to be senescence-associated beta-galactosidase (SAbGAL) positive to different degrees, which does not mean real senescence is occurring in these cells. ONG41008 did not harm normal cells and elicited massive senescence in pMFBs without apoptosis. However, ONG41008 caused massive senescence as well as apoptosis in cancer cells. In other words, ONG41008 was capable of sensing intracellular molecular environments between normal cells, cancer cells, and pMFBs. This molecular recognition capability prompted us to explore how ONG4008 behaved on A549 (a human lung adenocarcinoma), PANC1(malignant human ductal adenocarcinoma), and mdr+PC3 (multidrug-resistant human prostate cancer). TP53, p21, and p16 were induced and/or nuclear relocated, suggesting that ONG41008 was recognized by these cells. ONG41008 drove A549 and PANC1 at G2/M phase arrest during 48 hrs, resulting in massive mitotic collapse. All cells died. Moreover, the cisplatin-resistant mdr+PC3 was also eliminated by ONG41008. An array of common components of apoptosis were activated, and especially, induction of Mcl1 was especially notable. These senolytics features were reported to oncogene-induced-senescence (OIS), in which the expression of over two activated oncogenes in the embryonic fibroblasts caused massive senescence and cell death as well. And the signature expression of Mcl1, an anti-apoptotic protein (a long form), was notable but two kinds of short forms are pro-apoptotic proteins. OIS was conducted *in vitro* cell culture models and whether or not the presence of OIS counterpart *in vivo* remains to be delineated.

Taken together, we discovered a synthetic polyphenol referred to as ONG41008 was both senogenic and senolytic and its senescent impacts may make the cell cycles of the ONG41008-treated cancer cells immensely arrested at the G2/M phase, leading to mitotic slippage and cell death. This interesting observation may be able to create an idealistic anti-cancer modality, specifically killing cancer cells, but normal cells remain unharmed.

## Introduction

Cellular senescence, senescence, in short, is a primary universal program for cell protection and an irreversible trans-differentiation (1). We generated a synthetic polyphenol called ONG41008. ONG41008 rapidly induced senescence of pathologic myofibroblasts (pMFBs) not normal fibroblasts with profound specificity (2). We postulated ONG41008 *per se* may be able to differently sense intracellular molecular environments between in DHLF (diseased human lung fibroblasts from IPF patients) and in NHLF (normal human lung fibroblasts) due to significant differences in cellular homeostasis.

Flavones are members of the polyphenol family (3), a group of over 8,000 compounds that have been exclusively found in the plant kingdom (4). In general, these phytochemicals protect plants from radiation damage (5). Due to their anti-oxidant and anti-inflammatory potential, flavones have long been used to treat inflammatory diseases such as arthritis and asthma (6). As shown in Figure 1A, Chromone, 1,4-benzopyrone-4-one, is a central chemical scaffold from now on called chromone-scaffold (CS) constituting flavones, flavanols, and isoflavones (7), and the CS derivatives give rise to a diverse family based on branching out chemical residues coupled to the core CS framework (8). We previously confirmed while ONG41008 was both senogenic and senolytic, Dasatinib, Quercetin and Fisetin were only senolytic (2). It has been recently reported that two polyphenols, Luteolin and Procyanidin C1, forming a trimer, were also senolytic (9). While senescence-inducing compounds are very rare, senolytic drugs, collectively termed senolytics, have been discovered abundantly in the plant kingdom and specialized chemical library (over 80 senolytics) (10)

**Figure 1.**
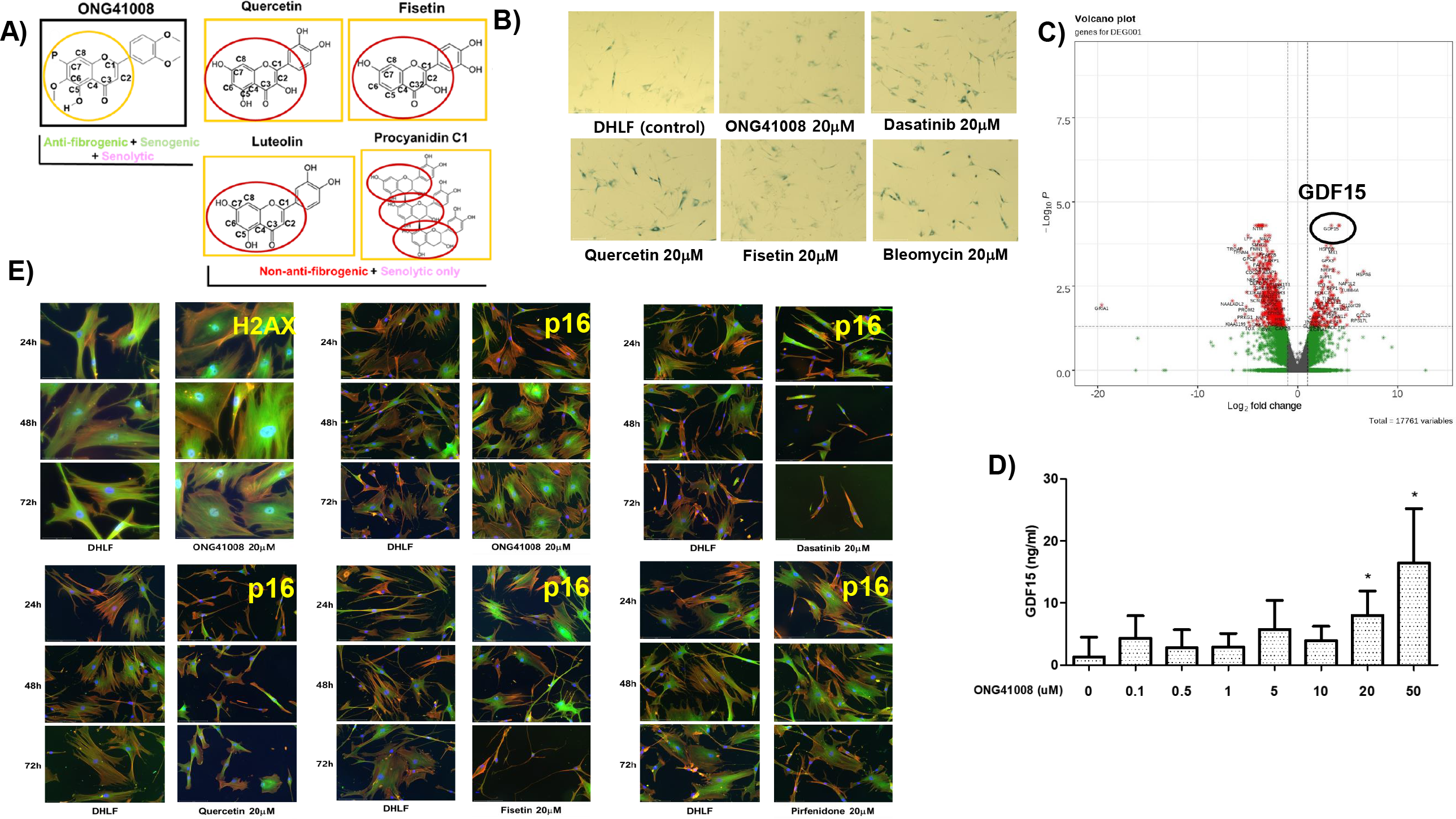
ONG41008 rapidly induced senescence in pathologic human pMFBs (DHLF) by ONG4108 and produced GDF15. **A**) Schematic representation of the polyphenols exhibiting senogenicity or senolysis. **B**) DHLF were stimulated with each 20μM of ONG41008, Dasatinib, Quercetin, Fisetin, or Bleomycin for 72 hrs, and fixed. SAβGAL assay was conducted. **C and D**) The DHLF-ONG41108 interactome axis was established via RNA-Seq. A volcano plot was derived from the DHLF-ONG41008 interactome. Soluble GDF15 was subjected to a GDF15 ELISA. **E**) DHLF were stimulated with 20μM of ONG41008, Dasatinib, Quercetin, or Fisetin for 24 hr through 72hrs. ICCs were conducted with the use of anti-H2XA and/or anti-p21.

Taken together, the senogenicity associated with ONG41008 as a polyphenol should be appreciated. In addition, ONG41008 also harbors a senolytic capability such that the combinatorial capabilities associated with ONG41008 can give rise to an idealistic anti-cancer modality which can distinguish cancer cells from normal cells.

## Results and Discussions

As shown in Figure 1B, DHLF, control cells, gave rise to pale green, suggesting that DHLF was already presenescent. Senescence-associated beta-galactosidase (SAβGAL) is thought to be a very early marker of senescence-associated-secretory-phenotype (SASP) (11). Upon stimulation of DHLF with ONG41008, Dasatinib, Quercetin, Fisetin, Bleomycin, or Pirfenidone, ONG41008, Dasatinib, and Fisetin modestly upregulated SAβGAL but ONG41008 only transformed DHLF into flattened morphology. Bleomycin showed a substantial SAβGAL upregulation, however, its cell morphology was unchanging, indicating that the production of SAβGAL is not always necessarily required for the acquisition of senescence. Prolonged stimulation of DHLF with Dasatinib led to massive cell death. Dasatinib was primarily developed as a Src kinase inhibitor for chronic myeloid leukemia (CML)(12). Quercetin or Fisetin is related to the mitochondrial channel formation via the Bcl2 and/or Bcl-xL family (13) and are several kinase inhibitors as well (14). Pirfenidone showed no differences in p21 expression or nuclear relocation. The induction of senescence of DHLF by ONG41008 was accompanied by massive flattened cell formation and rapid induction of GDF15 without apoptosis.

We explored the changes in the genome-wide gene expression profile of ONG41008 on DHLF. An interactome before or after treatment of DHLF by ONG41008 was established via RNA-Seq and was defined as the DHLF-ONG41008 axis. We observed hormone GDF15 was profoundly induced at transcription levels and produced as being secretory hormone by ELISA as shown in Figure 1C and Figure 1D). GDF15 belongs to the TGFβ supergene family and its receptor is a heterodimeric protein as constituted in combination with glial-derived neurotrophic factor-family receptor α-like (GFRAL) as an endogenous receptor and the rearranged during transfection (RET) tyrosine kinase co-receptor being a signaling receptor for GFD15 (15). GDF15 is dramatically induced upon cell injury and is required for tissue tolerance. None of the immunomodulatory SASP including TNFα, CCL2, or IL-6 for 72 hrs after ONG41008 stimulation was observed. We stimulated DHLF with ONG41008, Dasatinib, Quercetin, Fisetin, or Pirfenidone known as the first-class-anti IPF drug, and senescence was observed by looking at the p16 induction and nuclear localization or the inducible expression of a senescence-specific histone, H2AX, by using immunocytochemistry (ICC). As shown in Figure 1E, a massive induction of H2AX was only generated by ONG41008, and early induction of p16 was detected in the DHLF stimulated with Dasatinib or Fisetin in 24 hr. Afterward, the treated cells became lean and were subjected to cell death. Neither Quercetin nor Pirfenidone affected p16. Thus, the ONG41008-mediated senescence features were associated with remarkable changes in flattered cell shapes and relocation to the nucleus of p16. We showed that continuous stimulation of DHLF with ONG41008 was unable to kill DHLF upon senescence. In order to explore how GDF15 affected senescent DHLF, we generated a few versions of shRNA-GDF15 and infected senescent DHLF with these viruses, no particular features were manifested.

Taken together, we found ONG41008 was able to induce rapid and massive senescence in DHLF (pMFBs) without apoptosis, and GDF15 seemed a pioneering SASP derived from the senescence of pMFBs.

A549 were stimulated with ONG41008, Dasatinib, or Adriamycin (Doxorubicin) as shown in Figure 2A, control A549 already massively expressed SAβGAL, supporting again that cancer cells may be the most presenescent. ONG41008 upregulated SAβGAL at 10μM and flattened cell morphology was seen and remained unchanging at 20μM. Upon treatment of A549 with Dasatinib, massive cell death occurred. As we previously observed, the caspase 3/7 assay yielded high activities (2), suggesting apoptosis was likely due to DDR. Adriamycin was the first chemotherapeutic compound to shrink the sizes of solid tumors and was known to be a strong senescence inducer at early administration. With longer administration, cancer cells apparently died of DDR, and normal cells could be “by standing cells” to be harmed by Adriamycin (16). Adriamycin dramatically changed the cell morphology being enlarged A549 and kept dying the cells. Since ONG41008 was not lesser toxic to A549 in comparison to Dasatinib or Adriamycin, number of the senescent cells began to surge within 48 hrs during which the ONG41008-mediated senescence seemed to be complete and apoptosis was followed. We set up another set of caspases 3,7 assays with the use of ONG41008, Quercetin, or Fisetin. Since Quercetin and Fisetin were senolytics we hypothesized upon treatment either treatment did not lead to apoptosis. As shown in Figure 2B, Quercetin or Fisetin alone could not kill A549 and PANC1 probably due to lack of senescence degree, and the treated cells were likely dead through necrosis-like apoptosis, indicating Quercetin or Fisetin is naturally toxic to A549. The toxicities shown by these polyphenols shall potentially be a problem for dose escalation. On the contrary, the ONG41008-treated A549 started apoptosis at 10μM after 24 or 48hrs. This convinced us that ONG41008 was both senogenic and senolytic. We found ONG41008 eliminated a variety of primary lung cancer cell lines; H358: bronchiolar carcinoma (NSCLC), HCC366: squamous carcinoma (NSCLC), NH1299 large cell carcinoma (NSCLC), and H1650 adenocarcinoma (NSCLC).

**Figure 2.**
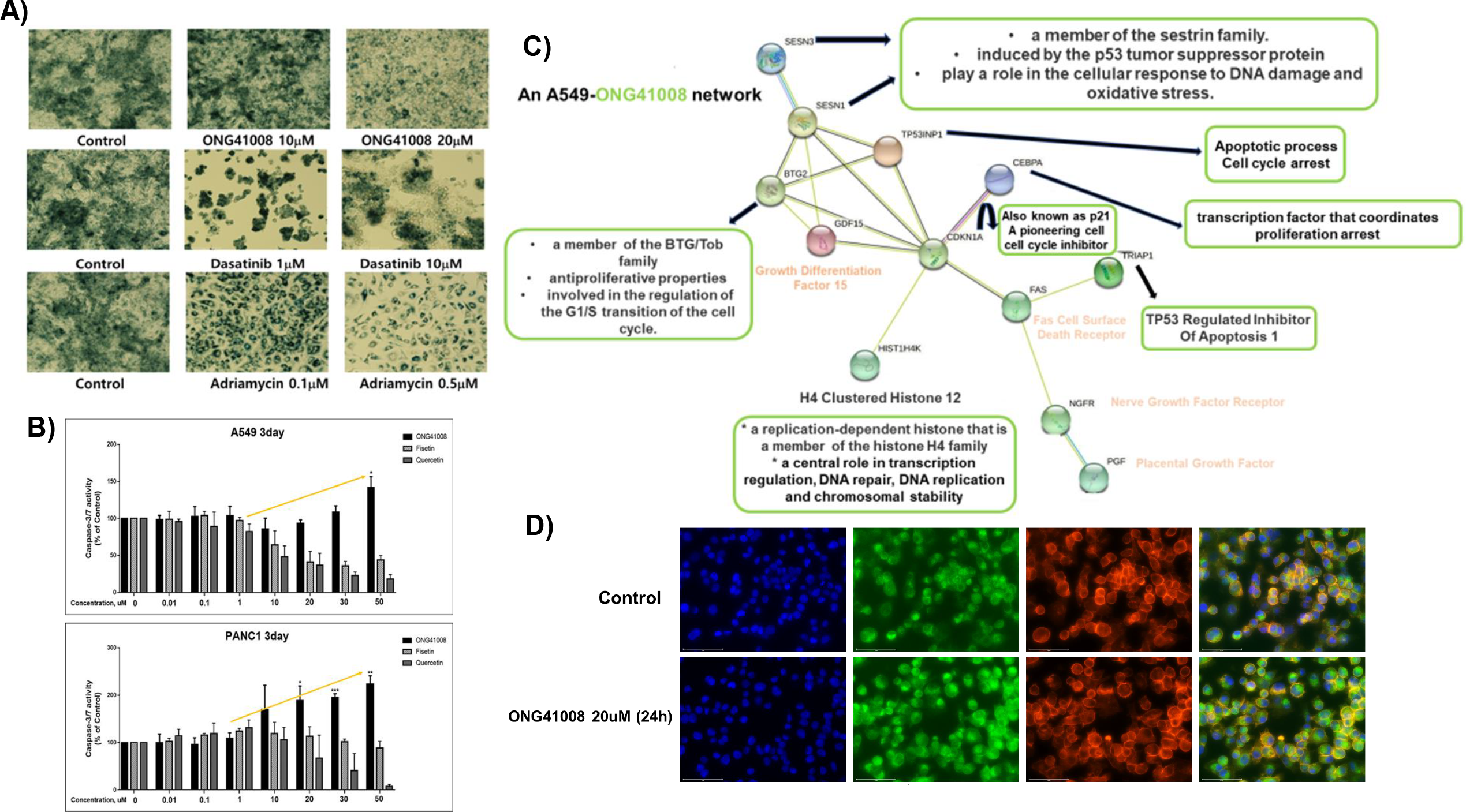
Genetic comparison, senolysis, and induction of Mcl1 associated with ONG41008. A549 cells were stimulated with designated drugs and an RNA-Seq was performed. And the A549-ONG41008 interactome was established. **A**) In order to confirm the existing expression of SAβGAS before or after stimulation with ONG41008, Dasatinib, or Adriamycin A549 at predetermined concentrations giving rise to reservation of color and morphological changes. was subjected to SAβGAL assays **B**) A549 or PANC1 were stimulated with ONG41008, Quercetin, or Fisetin, and Caspase 2,7 assays were conducted **C**) An A549-ONG41008 interactome was driven by an RNA-Seq. By using String program, interacting proteome were linked, and potential functions of each node were linked, giving rises to two hubs; 1) Apoptosis and 2) cell cycle regulation. **D**) Induction of Mcl1 by ONG41008; A549 were stimulated with 20μM ONG41008 for 24 hr. ICC was performed with the use of anti-Mcl1

An A549-ONG41008 interactome was established as seen in Figure 2C. We were astonished to discover CDKN1A (p21) was a central nod linked to GDF15, which may play an important role in the ONG41008-mediated senescence and senolysis. GDF15 has been known to be a potent apoptotic hormone in some neuronal cells (17). Moreover, there is a direct connection of FAS, TRIAP1, or nerve growth factor receptor (NGFR). NGFR negates the tumor suppressor p53 as a feedback regulator convinced us that this interactome was largely responsible for the apoptosis of A549 stimulated with ONG41008. The remaining proteins linked to p21 are BTG2 and CEBPA which have been implicated in cell cycle arrest.

Taken together, the A549-ONG41008 interactome suggests p21 plays a central role in the ONG41008-mediated senolysis. The hub located above p21 may be responsible for cell cycle arrest, antiproliferation of A549, and the quality control of DNA replication whereas the hubs below p21 is largely related to senolysis. Cancer cells were old cells. This extraordinary feature may happen due to their uncontrolled cell division, accumulation of DNA mutations, weakness of DDR capabilities, or oncogenic activations. Original senescence was found at the activation of H-ras. Thereafter, it has been appreciated that the addition of the second oncogenic mutation to H-ras led to massive senescence and drove the cell cycle to G2/M, thereby giving rise to oncogene-induced-senescence called OIS. OIS was typified by remarkable G2/M cell cycle arrest, leading to mitotic slippage or collapse. Massive cell death is now defined by senolysis. Although several apoptotic components have been found in common, senolysis was different from DDR-mediated apoptosis. Senolysis was exclusively dependent on senescence. Induction of Mcl1 was notable and this was contrasted to the original function, anti-apoptosis, of the full-length Mcl1. OIS held true for induction of Mcl1. We only speculate the short forms of Mcl1was massively induced and acted as apoptotic engager. This line of observations suggests that 1) ONG41008 can offer an idealistic anti-cancer modality, 2) cancer cells were more prevailingly presenescent as we saw A549 stimulated with ONG41008, and 3) the following senolytic capabilities would be lethal to A549 mainly due to G2/M mitotic collapse in conjunction with induction of Mcl1.

To prove this interesting hypothesis, we stimulated A549 or a primary NSCLC (H358) with ONG41008 for 72 hrs. As shown in Figure 3A, massive senescence occurred in A549, giving rise to flattened morphology as well as multinucleation denoted by white circles. Moreover, the ONG41008 drove H368 to reveal multinucleated cell colonies. A set of caspase3,7 assay clearly showed senolysis. These data strongly indicated ONG41008 was an idealistic armament killing cancer cells via its combinatorial therapeutic efficacies of senescence and senolysis. No normal cells are harmed. In addition, we performed intertumoral injection of A549 into in BALB/c for 60 days, establishing A549-xenografted tumors (Figure 3B). Tumor mass and tumor volume were remarkable regressed in the mice treated with ONG41008. Within 10μM to 50μM ONG41008 significant tumor volume and masses began to be decreased.

**Figure 3.**
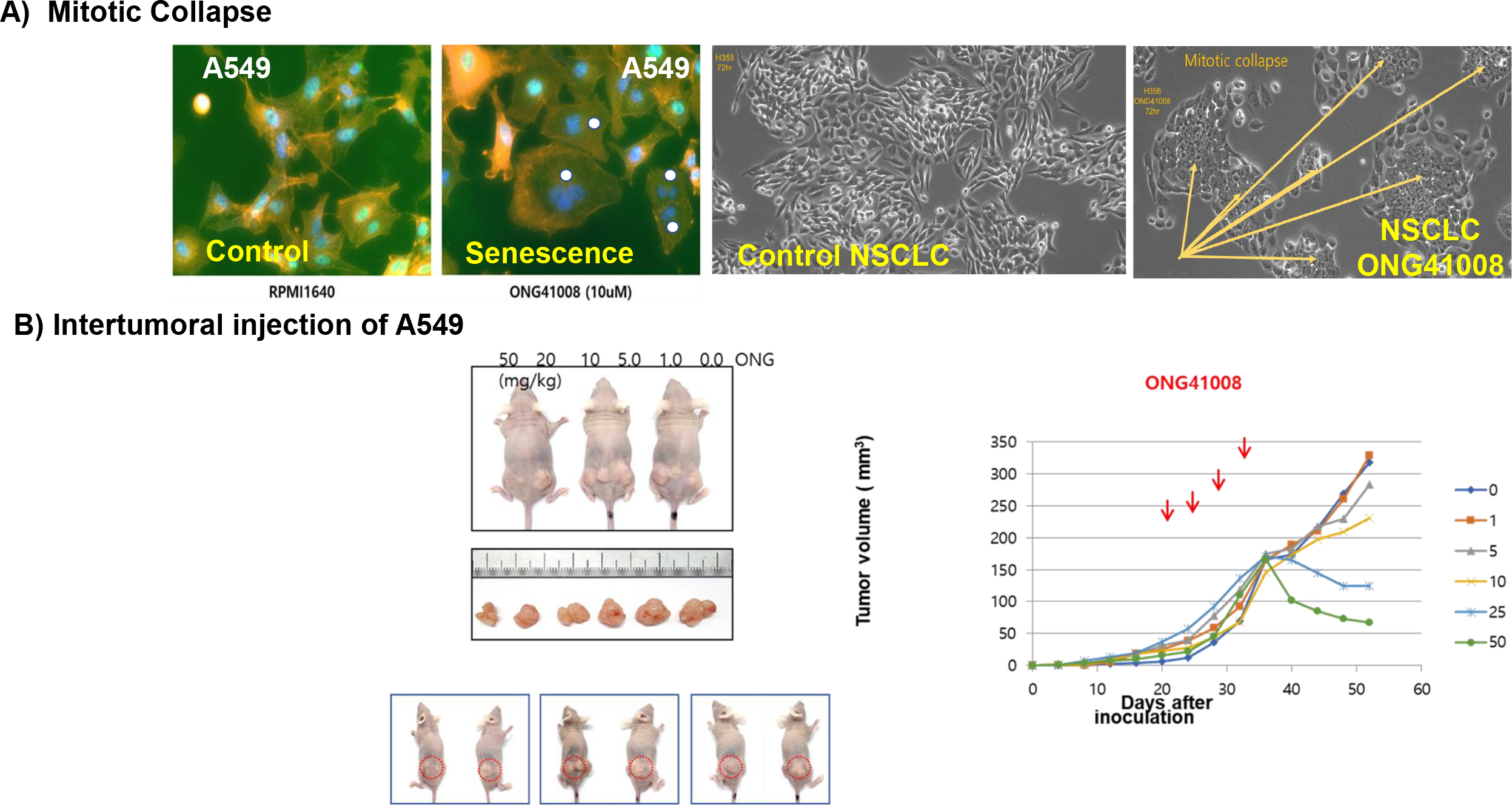
Induction of mitotic collapse and regression of tumors formed by A549-xenografeted in BALb/C. **A**) A549 or a primary NSCLC (H358) were stimulated by ONG41008 for 72hrs. Massive induction of A549 or mitotic collapse in conjunction with multinucleation was visualized. **B**) Formation the A549-xenografted tumor and therapeutic effects of ONG41008; Each 10^6^ of A549 cells were implanted into limbs, and a variety of ONG41008 concentrations dissolved in Geilcire 44/14 up to 50 mpk. After 60 days tumor masses were extracted, tumor volume and mass were measured.

In conclusion, ONG41008 was both senogenic and senolytic. ONG41008-mediated senescence and senolysis resembled OIS. The secondary viral oncogene infection to the H-ras expressing embryonic fibroblasts rapidly underwent massive senescence followed by apoptosis, and the induction of Mcl1 was notable (18). At this point, we cannot construct a full frame enabling us to reconcile or contrast to OIS as compared to the impact of ONG41008 to cancer cells. ONG41008 is a lipophilic compound. We found ONG41008 produced IP3 and DAG, suggesting that PI3K or PLC-γ got activated with ONG41008. It has been well known that these two lipid kinases are required for initiation and perpetuation of senescence. We do not know where ONG41008 was preferentially located at the intracellular compartments. But, upon stimulation of DHLF (pMFBs) with ONG41008 a rapid induction of senescence but reserved apoptosis. The degree of the senescence degree in DHLF stimulated with ONG41008 may not be sufficient for coupling to senolysis. We would like to call “senescence critical points (SCP)”, which would be diversified in all different cells. However, in the case of A549, primary NSCLC lines and different tissues-derived cancers were supposed to be significantly presenescent and ONG41008-generating capabilities of senescence were added to such that this combinatorial synergy could induce massive senescence and exerted to initiate senolysis including mitotic collapse. We demonstrated that the treatments of A549 with ONG41008 rapidly consistently changed in the levels of NAD to normal ranges. NAD of A549 per se was zero or minus (2). We could not explain how NAD in A549 augmented. In a good number of literatures, increase in NAD in cancer cells was deeply was related to reactive oxygen species (ROS), resulting in cancer cell death. Likewise, some results were oppositely described. This NAD experiments with the use of ONG41008 remains to be further explored. In addition, intracellular pH, oxygen concentrations, or ATP concentrations should be precisely measured upon a variety of ONG41008 concentrations. Lastly, our discovery called Cancer Cell-Specific Killing (CCSK) or Cancer Cell Senotherapy (CCS) can be an idealistic anti-cancer modality. Further translational studies should be conducted.

## Acknowledgements

This preprint has been indebted to MK Moon and S.B Kim for research assistances. Kind provision of human malignant cell lines should be appreciated to Dr. IW-Kim

## Grants and Outsourcing

These works were conducted possible due to provision of intramural investment from Osteoneurogen. And a part of the research funds was utilized to be contracted for animal experimentations with Dr. JI Gil working for the Korea University at an accredited germ-free animal husbandry.

## Materials and Methods

### Cell Culture, Reagents, and assays

DHLFs were purchased from Lonza (Basel, Switzerland) and cultured in fibroblast growth medium (FBM, Lonza, Walkersville, MD, USA). Recombinant human TGFβ and PDGF were obtained from Peprotech (Rocky Hill, CT, USA) and used at a final concentration of 5 ng/ml. Chemically synthesized ONG41008 was obtained from Syngene International Ltd. (Bangalore, India), dissolved at a stock concentration of 50 mM in DMSO, and stored in aliquots at −20 °C. DMSO was used as a control. The RAW264.7 cell line was purchased from the Korean Cell Line Bank (Seoul, Korea) and cultured in RPMI supplemented with 10% FBS and 1% P/S (Welgene, Seoul, Korea). LPS was purchased from Sigma and used at a final concentration of 100 ng/mL. Caspase-3 Assay The caspase-3 activity was measured using a caspase-3 assay kit (Abcam, ab37401) following the manufacturer’s protocol. Cells treated with various concentrations of ONG41008, pirfenidone, or nintedanib were harvested and lysed on ice. The protein concentration was then measured by a BCA assay (ThermoFisher, 23227) and adjusted to 50 μg protein per 50 μL cell lysis buffer. 4.7. Mitochondrial Membrane Potential Assay

### Immunocytochemistry and Western blotting

Cells were fixed using 4% paraformaldehyde, permeabilized with 0.4% TritonX100, blocked with 1% BSA, and incubated with rhodamine phalloidin (Thermo Fisher, Waltham, MA, USA), anti-GATA6 (Abcam, Cambridge, UK), anti-p53 (Cell Signaling Technology, Beverly, MA, USA), p21(Abcam, Cambridge, UK), p16-INK4A (Proteintech, Rosemont, IL, USA), and ZEB1 (Cell Signaling Technology, Beverly, MA, USA) for 4 h at room temperature. After washing, cells were incubated with an Alexa Fluor 488 (Abcam, Cambridge, UK)-conjugated secondary antibody. Images were analyzed using EVOS M7000 (Invitrogen, Waltham, MA, USA). The A549 cells were seeded at 1 × 10^6^ cells/well in 100 mm cell culture dishes and incubated overnight, followed by treatment with various concentrations of ONG41008. After 24 h, the cell lysates were clarified by centrifugation at 14,000× g for 10 min and the supernatant was collected. The protein concentrations were quantified by the Bradford assay (Thermo Fisher, Waltham, MA, USA). Thereafter, 25 μg of cellular protein was loaded on a 10% SDS-PAGE gel and transferred to nitrocellulose membranes. After blocking with 5% BSA, the membranes were incubated with anti-p53 (Cell Signaling Technology, Beverly, MA, USA), phosphor-p53 (Cell Signaling Technology, Beverly, MA, USA), p21 (Abcam, Cambridge, UK), p16-INK4A (Proteintech, Rosemont, IL, USA), and GAPDH (Abcam, Cambridge, UK) overnight at 4 °C. After washing thoroughly, membranes were incubated with an HRP-conjugated secondary antibody. Protein bands were visualized using ECL reagent (Abfrontier, Seoul, Korea) and Uvitec HD9 (Uvitec, Cambridge, UK).

### Live Imaging

DHLFs were seeded in 12-well cell culture plates and, after 24 h, were treated with TGF beta (5 ng/mL), nintedanib (10 μM), pirfenidone (10 μM), and ONG41008 (10 μM). Cells were incubated in an EVOS M7000 CO2 incubation chamber (Invitrogen, Waltham, MA, USA); cell morphology images were captured every 30 min.

### RNA-Seq, expression signature and interactome establishment

Differential Gene Expression, and Interactome Analyses Processed reads were mapped to the Mus musculus reference genome (Ensembl 77) using Tophat and Cufflink with default parameters. Differential analysis was performed using Cuffdiff using default parameters. Then, FPKM values from Cuffdiff were normalized and quantitated using the R Package tag count comparison (TCC) to determine statistical significance (e.g., p-values) and differential expression (e.g., fold-changes). Gene expression values were plotted in various ways (i.e., scatter, MA, and volcano plots) using fold-change values and an R-script developed in-house. The protein interaction transfer procedure was performed using the STRING database with the differentially expressed genes. A 60 Gb sequence was generated, and 10,020 transcripts were read and compared. The highest confidence interaction score (0.9) was applied from the Mus musculus reference genome, and information regarding interactions was obtained based on text mining, experiments, and databases (http://www.string-db.org/accessed on 6 December 2021). Due to the proprietary company information, we have not provided a detailed interpretation of the RNA-Seq or interactome data.

#### Real-Time PCR

Reverse Transcriptase PCR and Real-Time PCR Cells cultured in either 12- or 24-well plates were washed twice with cold PBS and harvested using a TaKaRa MiniBEST universal RNA extraction kit (Takara, Japan). RNA was purified using the same kit according to the manufacturer’s protocol. RNA was reverse-transcribed using a cDNA synthesis kit (PCRBio Systems, London, UK). Synthesized cDNA was amplified with StepOne Plus (Applied Biosystems, Life Technologies, Waltham, MA, USA) and 2× qPCRBio probe mix Hi-ROX (PCRBio, London, UK). Comparisons between mRNA levels were performed using the ΔΔCt method, with GAPDH as the internal control.

